# Alkaline and loess grasslands with contrasting richness and biomass patterns are not separated on the CSR strategy spectrum

**DOI:** 10.1101/2023.10.10.561704

**Authors:** Andrea McIntosh-Buday, Judit Sonkoly, Attila Molnár, Katalin Szél-Tóth, Viktória Törő-Szijgyártó, Szilvia Madar, Evelin Károlyi, Patricia Elizabeth Díaz Cando, Gergely Kovacsics-Vári, Béla Tóthmérész, Péter Török

## Abstract

Studying the relationship between biodiversity patterns and processes in vegetation has been at the centre of interest in vegetation ecology for several decades. By studying the biomass of loess and alkaline grasslands along a water and salinity gradient, we aimed to analyse species diversity and CSR functional type patterns. We aimed to test the following hypotheses: i) The biomass and species richness scores and the species composition are significantly different between the sampled grassland community types. ii) The sampled communities are well separated based on the CSR strategy spectrum. iii) The amount of green biomass and litter are positively correlated with competitiveness and negatively with stress tolerance. The biomass and species richness scores and the species composition of the sampled communities along the sampled gradients were significantly different; the highest species richness, evenness and Shannon diversity values were found in loess grasslands. The highest level of litter accumulation was found in alkaline meadows. The communities were well separated in the ordinations but surprisingly, calculation of coordinates for CSR strategy types have not shown clear separation of the grassland community types. Based on the results we can conclude that the current CSR classification is too robust to delineate grassland communities in alkaline landscapes which are markedly different in composition. We point out that the CSR classification is based on the magnitude of stress, and it is not able to differentiate between the various forms of stress which might be responsible for the marked compositional differences. Therefore, developing an index focusing specifically on the functional characterisation of stressed communities and able to differentiate between different forms of stress (e.g., water and/or salinity stress) would be highly beneficial for future studies.

## 1. Introduction

Studying the relationship between biodiversity patterns and processes in vegetation has been at the centre of interest in vegetation ecology for several decades (Watt, 1947; White, 1979; van der Maarel, 1996; Helm et al., 2014). Besides the classical analysis of species diversity and composition, the increasingly popular functional approach using plant traits and/or providing various classifications for ecological plant strategies is now recognised as a powerful tool for explaining and understanding vegetation patterns and underlying processes at multiple scales (Díaz and Cabido, 2001; Díaz et al., 2007). While analysing single trait patterns can reveal important background mechanisms, when multiple processes drive the community assembly, they might affect different traits differently. Because of this, analysing multiple traits together can provide more accurate approximation of functions (de Bello et al., 2010). There are several ways to integrate single traits into a more complex analysis. One possibility is to use multi-trait indices such as the three components of functional diversity: functional richness, functional divergence, and functional evenness suggested by Mason et al. (2005). However, several criticisms have arisen regarding the application of multivariate indices, significantly constraining their explanatory power due to challenges in selecting the most appropriate traits, numerous missing values for traits that are not easy to measure, and correlations among the selected traits (Lefcheck et al., 2015).

Another opportunity is to use ecological strategy classification, out of which the CSR classification of Grime (1977) is the most established one. Grime identified stress and disturbance as the two most important drivers of plant community assembly: while stress prevents biomass production, disturbance is responsible for the elimination of produced biomass (Grime, 1977, 1988). Based on the various levels of stress and disturbance, three major plant strategy types can be identified: competitors (C), stress tolerators (S) and ruderals (R). Competitors are able to grow and uptake resources fast, produce large amounts of biomass and have in general large size and thus effectively compete with others for light and/or other resources. In contrast, stress tolerators display a conservative resource uptake and retention strategy, have a long lifespan, and grow slowly, but display a high tolerance to stress. Ruderals are adapted to environments characterised by high levels of disturbance and low levels of stress – they have high growth rate, effective and rapid resource uptake, and generally short lifespan and a rapid lifecycle. The CSR strategy system is frequently represented by a continuum rather than by discrete strategy types, and most plants display mixed strategies (Grime and Pierce, 2012). There are several methods of CSR classification developed mostly for herbaceous vegetation in Europe (Grime et al., 1988; Hodgson et al., 1999; Pierce et al., 2013), but the most recent classification by Pierce et al. (2013) was used for global analyses of the CSR spectrum and was validated to be a robust method comparing different regions and floras (Pierce et al., 2017). The other benefit of the Pierce et al. (2013) method is that it uses easily measurable leaf traits, for which a global dataset and several regional datasets are available for analyses (Kattge et al., 2020; Tavşanoğlu and Pausas, 2018; Sonkoly et al., 2023). The applicability of the classification was tested in several plant communities so far, but, to our knowledge, saline habitats were not included (Caccianiga et al., 2006; Cerabolini et al., 2010; Negreiros et al., 2014).

Several former studies analysed vegetation patterns and zonation in saline habitats (e.g., Suchrow and Jensen, 2010; Kelemen et al., 2013; Valkó et al. 2014; González-Alcaraz et al., 2014). Saline habitat complexes form excellent study objects as they have i) a rather limited species pool constituting relatively simple plant communities, and because ii) their vegetation zonation is driven, besides local topography and macroclimate, by a dynamic spatial and temporal relationship of salt contents and soil inundation (Piernik, 2012; Kelemen et al., 2013; Valkó et al., 2014; Matinzadeh et al., 2022). Studying vegetation gradients along increasing elevation, water availability and/or stress is important for understanding community dynamics and patchiness in alkaline landscapes. While coastal wetlands and inland salt marshes were the subjects of several previous studies, inland salt grasslands with alkaline soils received less attention (but see Kelemen et al., 2013 or Valkó et al., 2014).

Kelemen et al. (2013) studied the relationship between biomass and species richness within alkaline landscapes. Their study encompassed a variety of grassland types, such as loess grasslands, short-grass steppes, and alkaline meadows. The research findings validated that the biomass-species richness relationship in alkaline grasslands follows a humped-back (unimodal) curve. This pattern was observed in grasslands from different geographical locations, demonstrated by studies of Mittelbach et al. (2001) or Fraser et al. (2015). This finding underscores the existence of a consistent and generalizable ecological phenomenon across diverse grassland environments. Short-grass steppe plots characterised by low biomass and species richness were situated at the ascending section of the biomass-species richness curve. Plots of alkaline meadows are characterised by the highest biomass production associated with low species richness were situated at the descending section of the curve. The loess grasslands with an intermediate biomass production and high species richness were situated at the peak of the humped-back curve. In general, the increase in species richness at the ascending section of the curve is explained by several factors including the decreasing levels of stress, while the descending section is explained mostly by the increasing levels of competition (Kelemen et al., 2013; Michalet et al., 2006; Fraser et al., 2015). At the peak of the curve, balanced conditions of stress and competition can be expected (Kelemen et al., 2013).

By studying the biomass of loess and alkaline grasslands along a water and salinity gradient we aimed to analyse species diversity and CSR functional type patterns. We tested the following hypotheses: i) The biomass and species richness scores and the species composition are significantly different between the sampled grassland communities. ii) The sampled communities are well separated based on CSR strategy spectrum. We expected the highest degree of S-selection in the short-grass steppe plots, while the highest degree of ruderality and competitiveness in loess grasslands and alkaline meadows, respectively. We also expected that iii) the amount of green biomass and litter are positively correlated with competitiveness and negatively with stress tolerance, as highly productive communities are characterised by species with competitor strategies.

## 2. Materials and methods

### 2.1. Study region

The study area is located in the Zám puszta, which forms an integral part of the Hortobágy puszta in the Hortobágy National Park, East Hungary. Hortobágy puszta, with about 75 thousand hectares is among the largest continuous natural-and semi-natural grassland area in Europe. Its vegetation is characterised by alkaline and loess grasslands and alkaline marsh habitats (Török et al., 2011). The climate of the area is moderately continental with a mean annual temperature of 10.5-11.0 °C and with a mean annual precipitation of 500-550 mm. The yearly maximum of the precipitation is in June with 65-80 mm on average, but high year-to-year fluctuations are typical (Pécsi, 1989; Lukács et al., 2015). The puszta grasslands were formed well before the last glacial maximum (LGM) (Sümegi et al., 2013) with continuous alkaline character. Later, wild grazers have been replaced by domestic livestock and several local varieties of domestic grazers have been bred (e.g., Hungarian grey cattle, racka sheep). The alkaline habitats form a very patchy and dynamic vegetation mosaic influenced by the spatially and temporally uneven patterns of soil moisture and salinity (Kelemen et al., 2013; Valkó et al., 2014), only a few centimetres of elevation difference may drive the formation of a considerably different species composition.

### 2.2. Sampled grassland community types

Loess grassland vegetation (*Festucion rupicolae*) historically covered the highest elevations in the region where chernozem black soils have been formed (Borhidi et al., 2012). These chernozemic soils were the most appropriate for agricultural crop production, so loess grasslands survived only in places where the loess grassland patches were inaccessible for crop production (like in kurgans or road verges, Deák et al., 2023), or were small and surrounded by extent alkaline vegetation. Characteristic graminoids are *Festuca rupicola*, *F. valesiaca*, *Carex praecox, Koeleria cristata*, and *Poa angustifolia*. Loess grasslands are usually characterised by the high cover and richness of forbs with several characteristic loess species including *Centaurea scabiosa*, *Linaria biebersteinii, Salvia nemorosa*, *S. austriaca*, *Phlomis tuberosa*, *Veronica prostrata*, and *Thymus glabrescens*. Small fragments of these grasslands are typically abandoned or mown, stands embedded in alkaline grasslands are frequently grazed by cattle or sheep (Török et al., 2011, 2018; Deák et al., 2020).

Short-grass steppes (*Festucion pseudovinae*) are typically situated between the elevation range of loess grasslands and alkaline meadows and according to the salinity of their soil, they can be either *Artemisia* short-grass steppes or *Achillea* short-grass steppes, and these two grassland community types often form a dynamic mosaic. Both grassland community types can be characterised with a high cover of *Festuca pseudovina*. The more saline *Artemisia* alkali steppes are generally covered by a shallow water up to a few centimetres in the spring and can be characterised also by the high cover of *Artemisia santonicum*. Beside of these species several characteristic forb species occur in these grasslands like *Aster tripolium ssp. pannonicus*, *Cerastium dubium*, *Bupleurum tenuissimum*, *Gypsophila muralis*, *Limonium gmelinii*, *Lotus tenuis*, *Podospermum canum* and several *Trifolium* species including *T. retusum*, *T. angulatum* and *T. fragiferum*. *Achillea* alkali steppes are less saline and characterised by a high cover of *Achillea collina* and *A. setacea* and also by several forb species like *Plantago lanceolata*, *Ornithogalum kochii*, *Ranunculus pedatus*, *Trifolium striatum, T. retusum* and *T. strictum*. Heavily grazed stands of both community types are also characterised by a high cover of thistles (e.g., *Cirsium vulgare*, *Carduus acanthoides*, *C. nutans*), or *Ononis spinosa* and *Cynodon dactylon*. These community types of grasslands are managed by moderate to intensive grazing by sheep or cattle (Török et al., 2012).

Alkaline meadows (*Beckmannion eruciformis*) are typically situated in an intermediate position between short-grass steppe grasslands and low-lying alkali marshes. The vegetation is in general species poor and characterised by high cover of tall-grass species like *Alopecurus pratensis*, *Elymus repens*, *Beckmannia eruciformis*, and *Agrostis stolonifera*, and tall forb species like *Leonurus marrubiastrum*, *Lycopus europaeus*, *Lythrum virgatum*, *Rumex crispus* and *R. stenophyllus*. Typical small forb species are also present in a second vegetation layer, the most frequent species are *Cerastium dubium*, *Galium palustre*, *Inula britannica*, *Lysimachia nummularia*, *Mentha aquatica*, *Ranunculus lateriflorus*, and *R. repens*). Also, several short rushes may appear in the vegetation (*Juncus compressus*, *Eleocharis* spp.). This habitat is generally covered by shallow water in the spring to early or late summer. The vegetation was traditionally maintained by extensive cattle grazing or typically after the Second World War by mowing for hay (Török et al., 2012; Deák et al., 2014).

### 2.3. Sampling setup

We selected three natural alkaline depressions in the surroundings of the village of Hortobágy (East Hungary, Figure 1). All of the depressions were imbedded into a natural alkaline landscape in the protected area of the Hortobágyi National Park. The sampled depressions were situated at least 500 m from each other. In May 2022, we sampled three grassland communities along transects starting at the highest elevations in loess grasslands, passing through short-grass steppes at intermediate elevations, finally reaching the low-lying alkaline meadows. Altogether three transects were sampled whose length were 30 to 50 m depending on the length of the vegetation gradient. Starting from the loess grasslands at the highest elevations, we sampled the biomass of the grasslands with a 20 cm × 20 cm wooden frame at 1 m intervals (Figure 1). We collected all aboveground biomass, including cryptogams (mosses, lichens, and large terrestrial algae), standing litter and the litter layer into paper bags. The harvested biomass was transported to the laboratory and dried in an exicator (65 °C, until weight constancy, but at least 72 hours). We sorted the biomass into main fractions, i.e., to litter (litter layer + standing litter), green biomass, and cryptogam biomass (not sorted further). Green biomass was further sorted into species. All biomass was weighed by a balance of 0.01 g accuracy. To support the biomass sorting, we made a list of species in a 1 ha area in the surroundings of the transects. Most communities were clearly separated already in the field; a few transitional plots were assigned to one of the bordering communities. In total, there were 30 biomass samples collected from loess grasslands, 45 from short-grass steppes, and 35 from alkaline meadows. The study sites were lightly to moderately grazed by beef cattle in the vegetation period.

**Figure 1.**
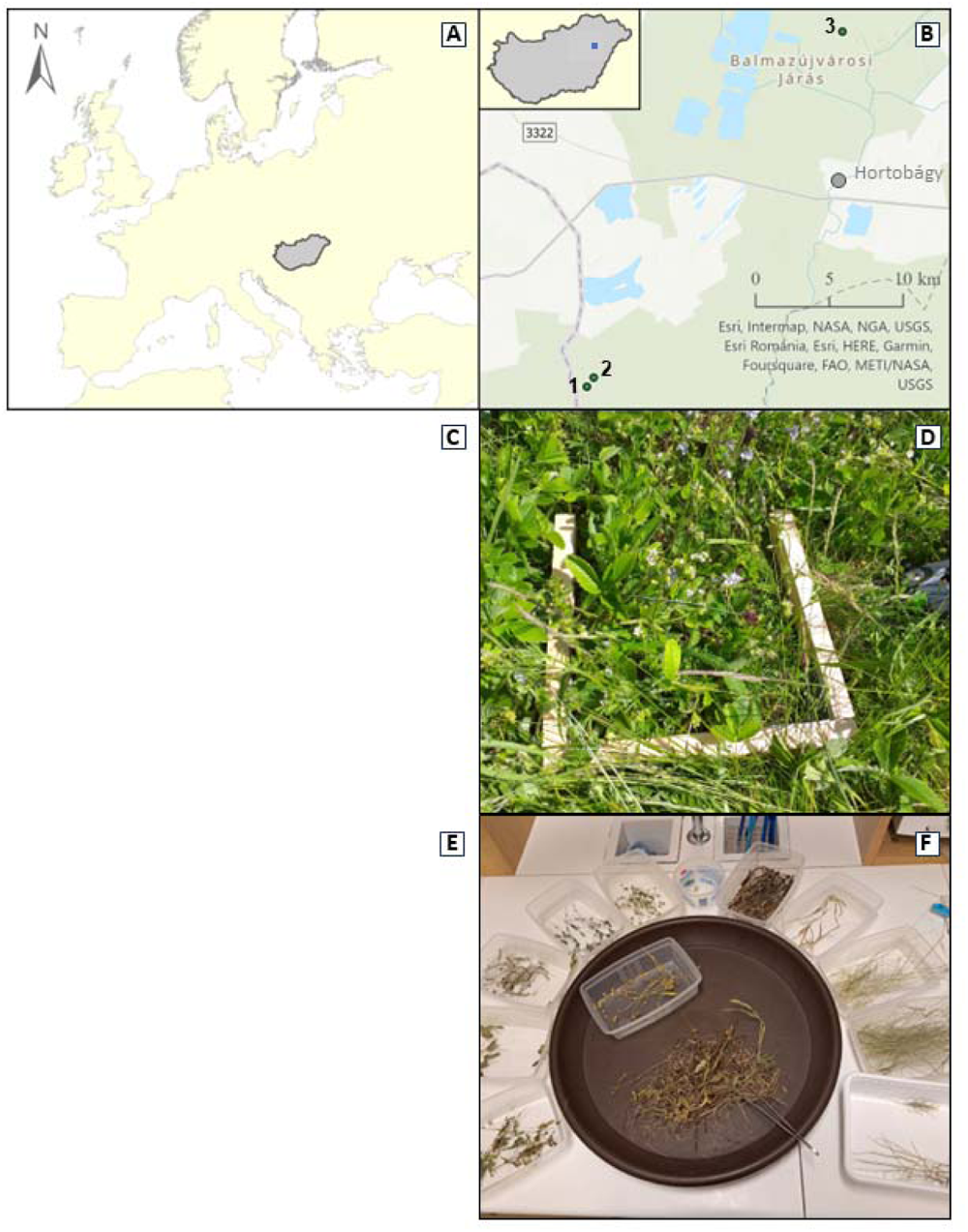
Location of study area (A, B; 1-3: our transects), field sampling (C, D) and biomass sorting (E, F). The figure section C and F was removed by the request of the bioRxiv as their policy is to avoid the inclusion of photographs and any other identifying information of people.

### 2.4. Data capture and analyses

To assign the species to CSR strategy types we used the CSR classification method of Pierce et al. (2013, 2017). This type of CSR strategy assignment is based on leaf traits such as leaf area (LA), leaf dry matter content (LDMC), and specific leaf area (SLA). The method is proven to be an effective way to classify species into various mixed CSR strategy types. It corresponds well with Grime’s theoretical CSR triangle and was proven to be sufficiently precise to distinguish the strategies of species within the same genera. As leaf trait data, we used our own measurements and other regionally measured data, both stored in the Pannonian Database of Plant Traits (PADAPT, Sonkoly et al., 2023). We used the FDiversity program package to calculate Shannon diversity (Shannon, 1948), Pielou’s evenness (Pielou, 1975), and community-weighted means (hereafter CWMs) of CSR coordinates (Casanoves et al., 2011). To evaluate the correlations between biomass fractions and CSR coordinates, we used the linear fit option of OriginPro 2018 (Pearson’s product moment correlation, OriginLab Corporation, 2017). To compare diversity, biomass, and CSR coordinates of the sampled grasslands, we used GLMMs where the fixed factor was community type, and the random factor was the transect identity. The GLMMs were calculated by SPSS 26.0 program package (IBM Corp., 2019). The species composition of the biomass of the sampled communities was compared with DCA ordination. We calculated separate ordinations based on presence-absence datasets and based on the species’ biomass. The ordinations were calculated by CANOCO 5.0 program package (Šmilauer and Lepš, 2014). Species nomenclature follows Király (2009).

## 3. Results

### 3.1. Species richness and biomass

The sampled communities were significantly different in terms of species richness, Shannon diversity, evenness and the mean weight of all biomass fractions except for cryptogam biomass (Table 1). The highest species richness, Shannon diversity and evenness were found in the loess grasslands, followed by short-grass steppes and alkaline meadows (Figure 2A-C). The green biomass was the lowest in short-grass steppes, while the litter was the highest in alkaline meadows (Figure 2D-E). We detected no significant differences between the three grassland community types in the biomass of cryptogams, but higher scores were typical in short-grass steppes (Figure 2F). The communities were well separated in the ordinations based on both presence-absence data and species abundance data; however, the point clouds of the loess grasslands and short-grass steppes had some minor overlap (Figure 3 and 4). The differences in species composition and biomass were also well reflected in the DCA ordination. In Figure 5, it is clearly shown that the increase in green biomass is pointed towards the loess grasslands, litter accumulation towards the alkaline meadows, while the increase in cryptogam biomass towards the short-grass steppes (Figure 5).

**Figure 2.**
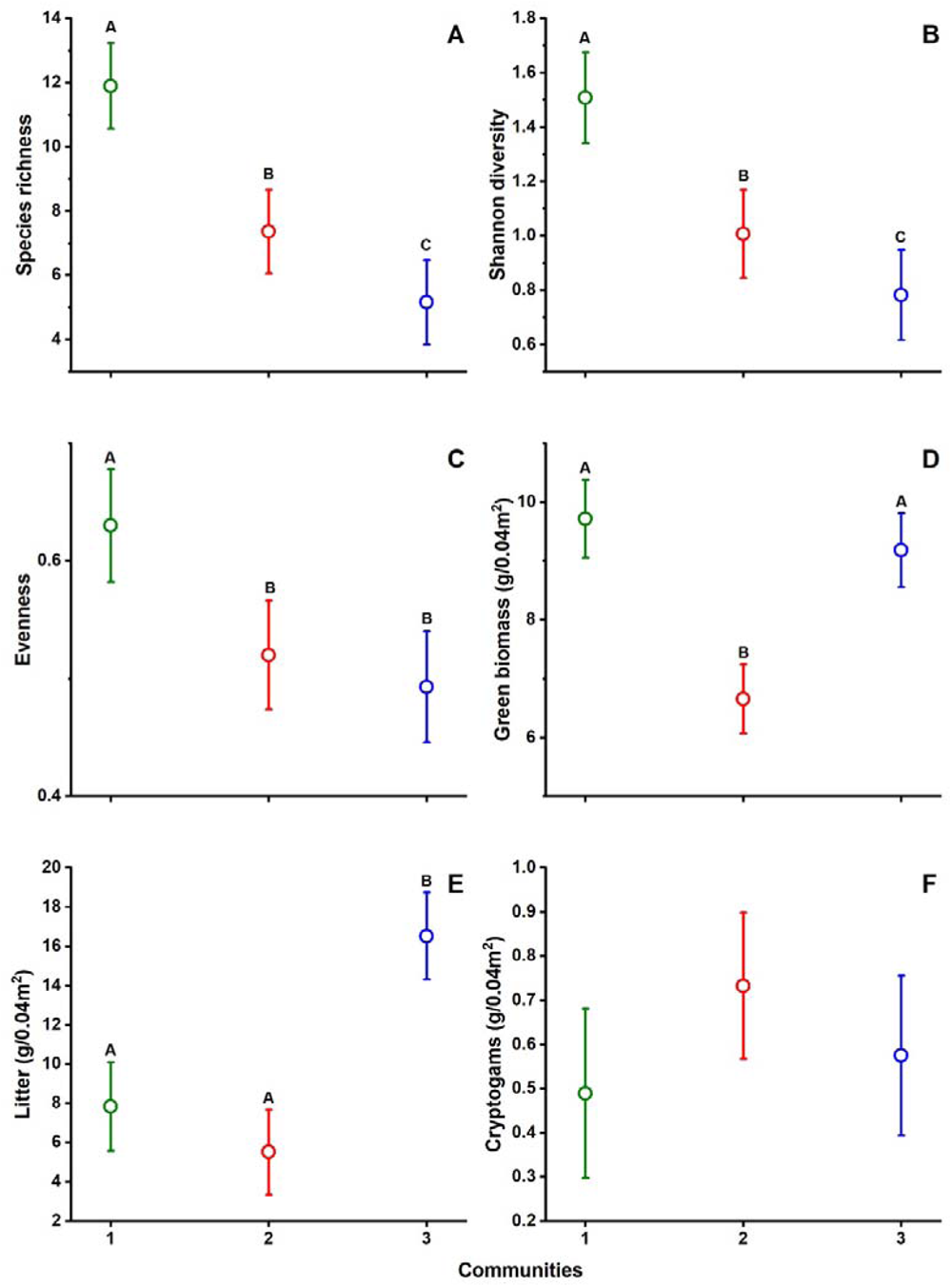
Community-weighted means of species diversity and the weight of various biomass fractions in the studied communities (mean ± SE, GLMM and least significant differences test, significant differences are indicated with different capital letters). Grassland community type codes: 1 = loess grasslands, 2 = short-grass steppes, 3 = alkaline meadows.

**Figure 3.**
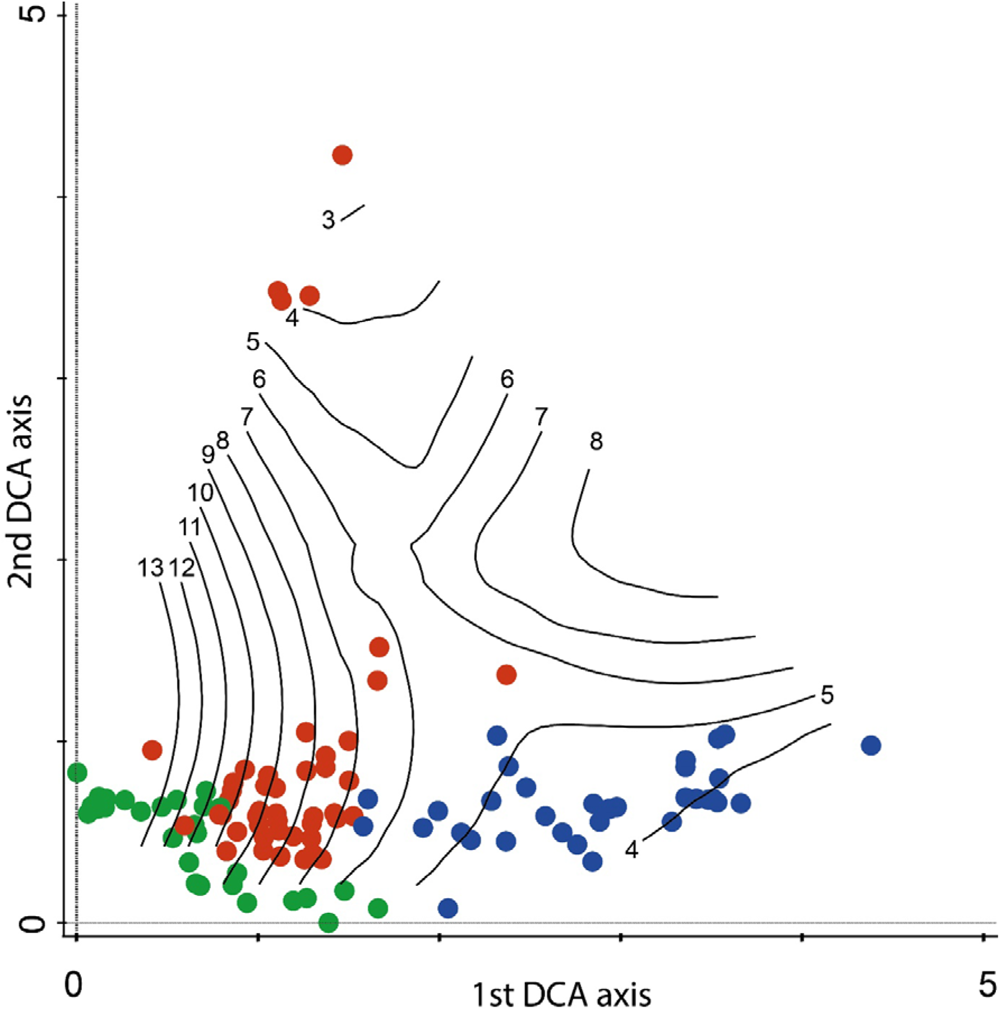
The species composition of the studied communities displayed by a DCA ordination (based on presence-absence data, species isolines are shown with black lines with species numbers). Grassland community type colours: green = loess grasslands, red = short-grass steppes, blue = alkaline meadows.

**Figure 4.**
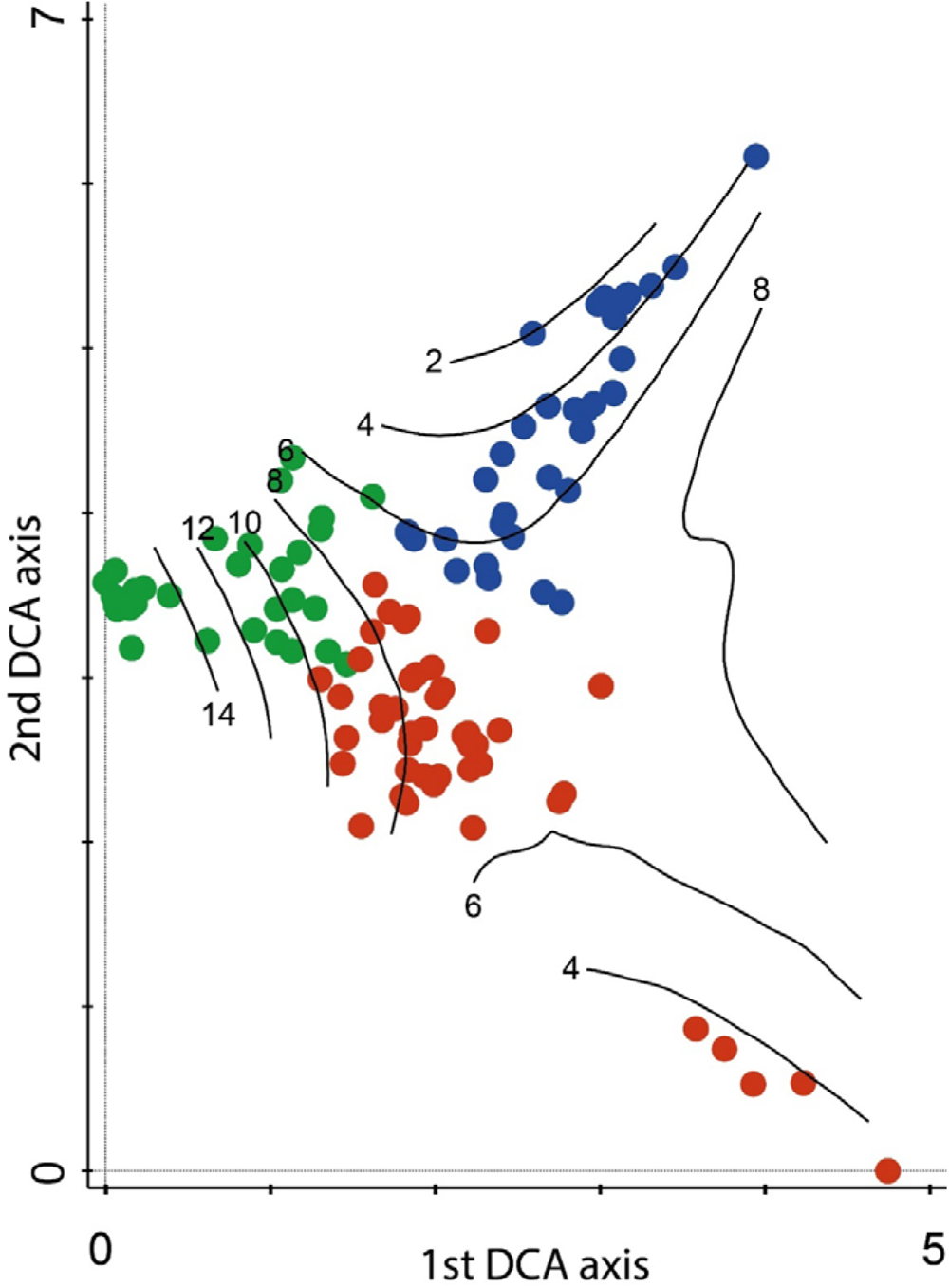
The species abundances of the studied communities displayed by a DCA ordination (species isolines are shown with black lines with species numbers). Grassland community type colours: green = loess grasslands, red = short-grass steppes, blue = alkaline meadows.

**Figure 5.**
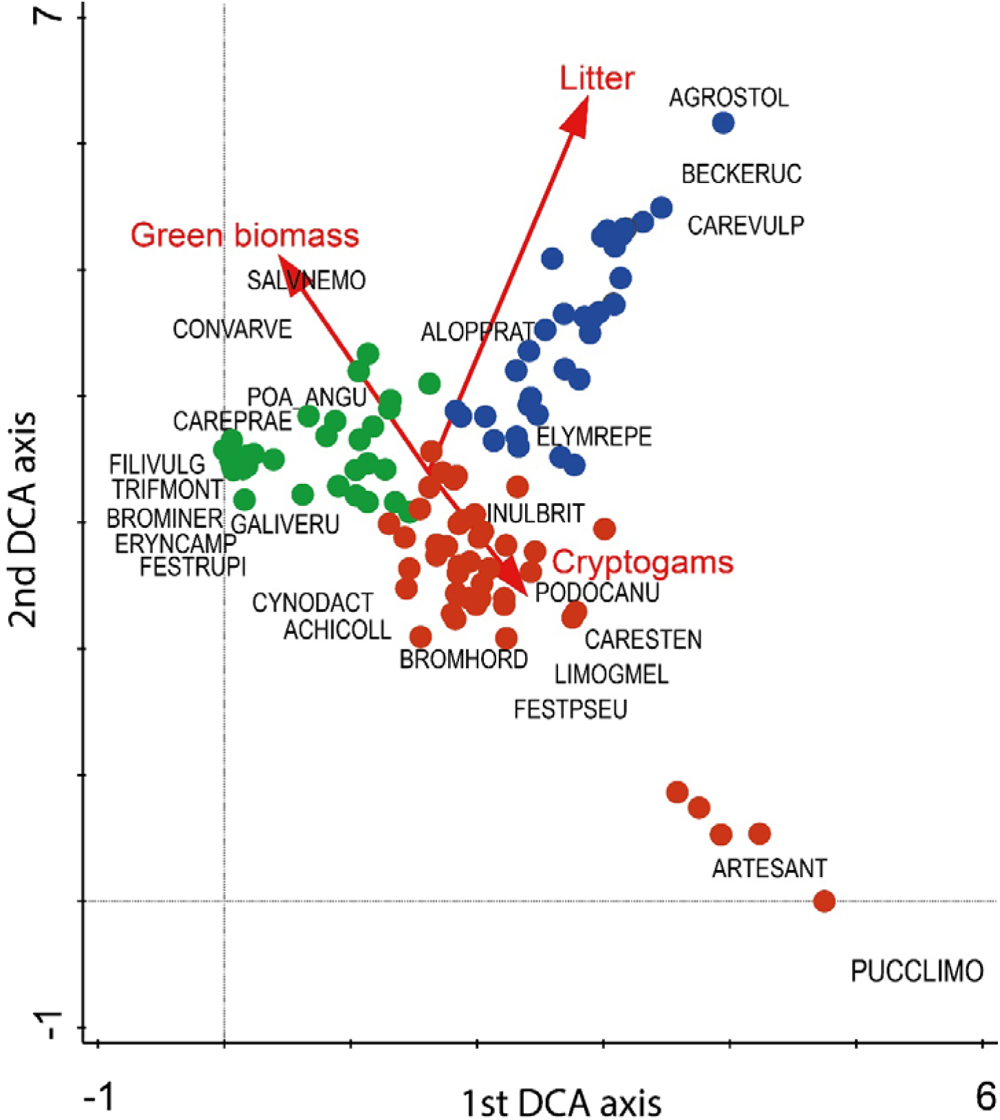
The relationship between species abundances (main dataset) and main biomass fractions (secondary overlay) of the studied communities displayed by a DCA ordination. Grassland community type colours: green = loess grasslands, red = short-grass steppes, blue = alkaline meadows. The most abundant 25 species are shown with an abbreviation of four letters of the genus and four letters of species names as follows in order of decreasing total biomass: ALOPPRAT: Alopecurus pratensis, FESTPSEU: *Festuca pseudovina*, ELYMREPE: *Elymus repens*, POA_ANGU: *Poa angustifolia*, CAREPRAE: *Carex praecox*, ACHICOLL: *Achillea collina*, FESTRUPI: *Festuca rupicola*, AGROSTOL: *Agrostis stolonifera*, ARTESANT: *Artemisia santonicum*, LIMOGMEL: *Limonium gmelinii*, TRIFMONT: *Trifolium montanum*, GALIVERU: *Galium verum*, SALVNEMO: *Salvia nemorosa*, ERYNCAMP: *Eryngium campestre*, FILIVULG: *Filipendula vulgaris*, CARESTEN: *Carex stenophylla*, CYNODACT: *Cynodon dactylon*, BROMINER: *Bromus inermis*, CONVARVE, *Convolvulus arvensis*, BECKERUC: *Beckmannia eruciformis*, CAREVULP: *Carex vulpina*, PODOCANU: *Podospermum canum*, BROMHORD: *Bromus hordeaceus*, PUCCLIMO: *Puccinellia limosa*, INULBRIT: *Inula britannica*.

**Table 1.** Effect of grassland community type on species diversity, CWMs of leaf traits and Pierce CSR coordinates. Significant effects are denoted with **bold face** (One-way GLMM, *p*<0.05, fixed factor – grassland community type, random factor – transect identity).

| Characteristic | Statistics |  |
| --- | --- | --- |
| | $F_{2,107}$ | $p$ |
| **Species diversity** |  |  |
| Species richness | **40.388** | **<0.001** |
| Shannon diversity | **33.533** | **<0.001** |
| Evenness | **7.596** | **0.001** |
| **Biomass** |  |  |
| Green | **10.429** | **<0.001** |
| Litter | **30.767** | **<0.001** |
| Cryptogams | 0.625 | 0.537 |
| **Leaf traits** |  |  |
| LDMC | **36.734** | **<0.001** |
| SLA | **4.213** | **0.017** |
| LA | **7.416** | **0.001** |
| **CSR coordinates** |  |  |
| Competitiveness | **11.901** | **<0.001** |
| Ruderality | **11.390** | **<0.001** |
| Stress tolerance | 0.234 | 0.792 |

### 3.2. Traits and ecological strategies

All of the community-weighted means (CWMs) of the studied leaf traits were significantly affected by the grassland community type (Table 1). The LDMC showed a clear and increasing gradient from loess grasslands to alkaline meadows (Figure 6A). The CWM of SLA was the highest in the short-grass steppes, while the CWM of LA was the highest in alkaline meadows (Figure 6B-C). Surprisingly, the analyses using weighted (Figure 7) and unweighted (Figure 8) calculation of coordinates for CSR strategy types have not shown clear separation of the communities. All communities were characterised by high levels of stress tolerance, moderate competitiveness and low ruderality, as expressed in the CWMs or S, C and R coordinates, respectively. However, there were significant differences between biomass-weighted C and R coordinates. The CWM value for C coordinates was the lowest, while that of R coordinates was the highest in short-grass steppes (Figure 6D-E). The stress tolerance of communities, expressed by the CWMs of S coordinates, were not significantly different between the communities (Figure 6F). We also calculated correlations between green biomass, litter, and the CSR coordinates. We found that C coordinates correlated positively both with the litter and green biomass (Figure 9A and D). The coordinates of S and R responded differently: while S coordinates correlated negatively with green biomass, there was no such relationship with litter (Figure 9B and E). R coordinates correlated negatively with the litter, but no such correlation was found with green biomass (Figure 9C and F).

**Figure 6.**
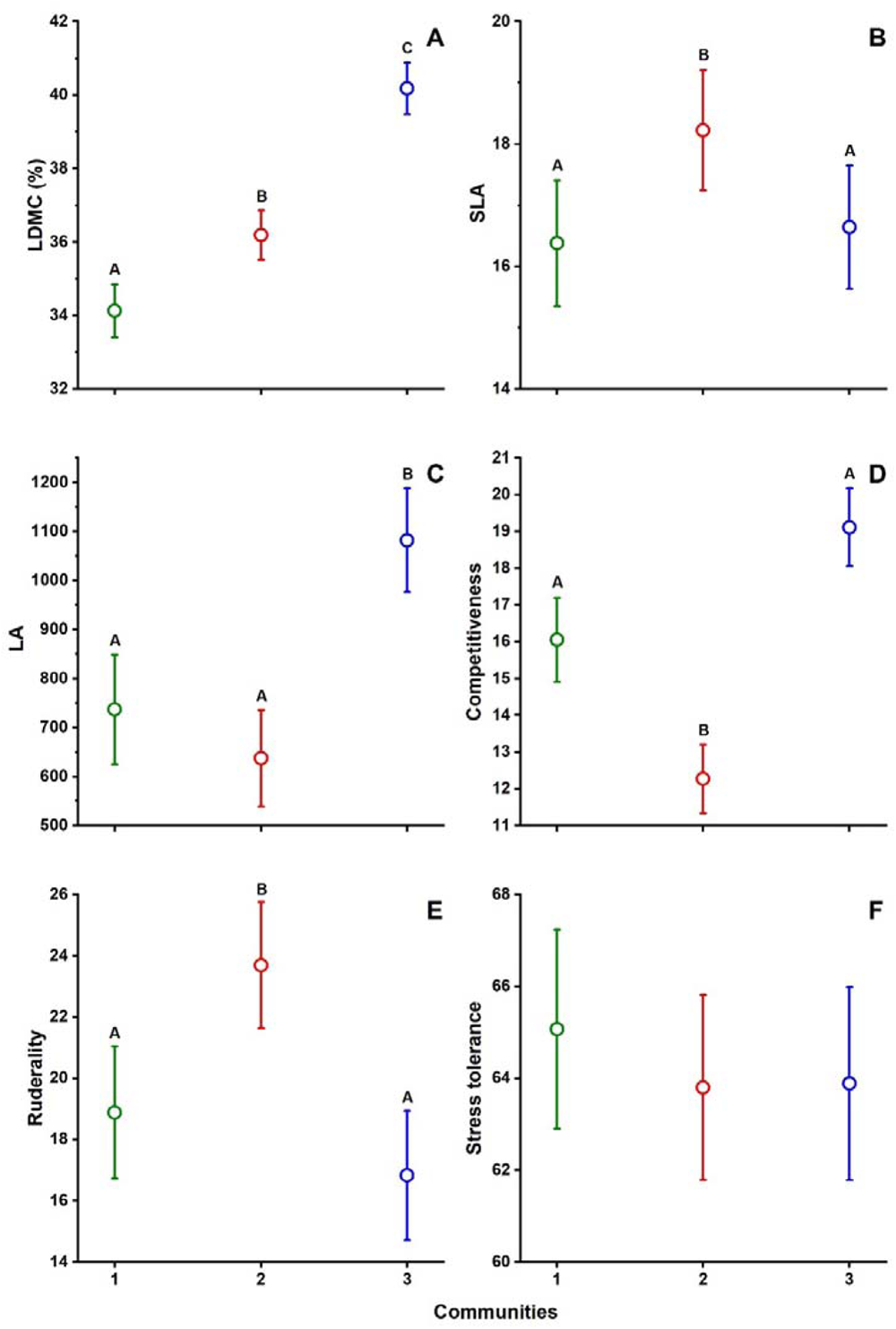
Community-weighted means of leaf traits, competitiveness, stress tolerance and ruderality expressed with the CWMs of CSR coordinates in the studied communities (mean ± SE, GLMM and least significant differences test, significant differences are indicated with different capital letters). Grassland community type codes: 1 = loess grasslands, 2 = short-grass steppes, 3 = alkaline meadows.

**Figure 7.**
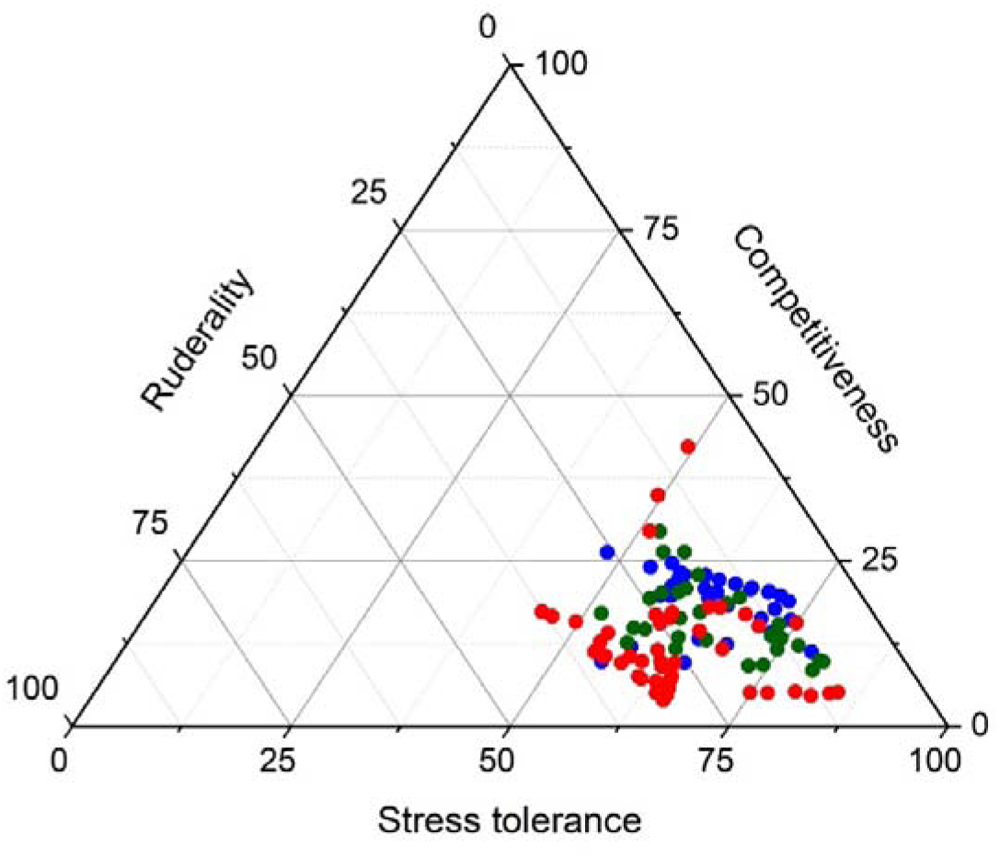
The community-weighted means of CSR strategy in the sampled plots, calculated by using the species’ biomass as weights. Grassland community type colours: green = loess grasslands, red = short-grass steppes, blue = alkaline meadows.

**Figure 8.**
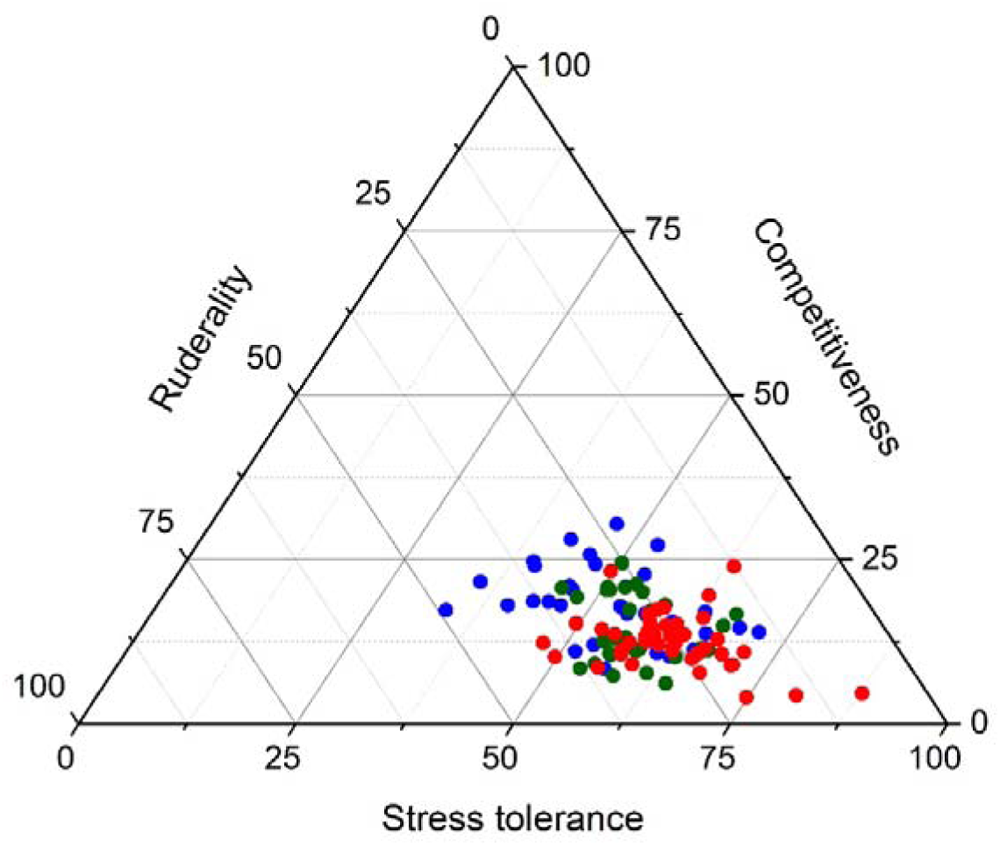
The means of CSR strategy calculated using un-weighted presence-absence data. Grassland community type colours: green = loess grasslands, red = short-grass steppes, blue = alkaline meadows.

**Figure 9.**
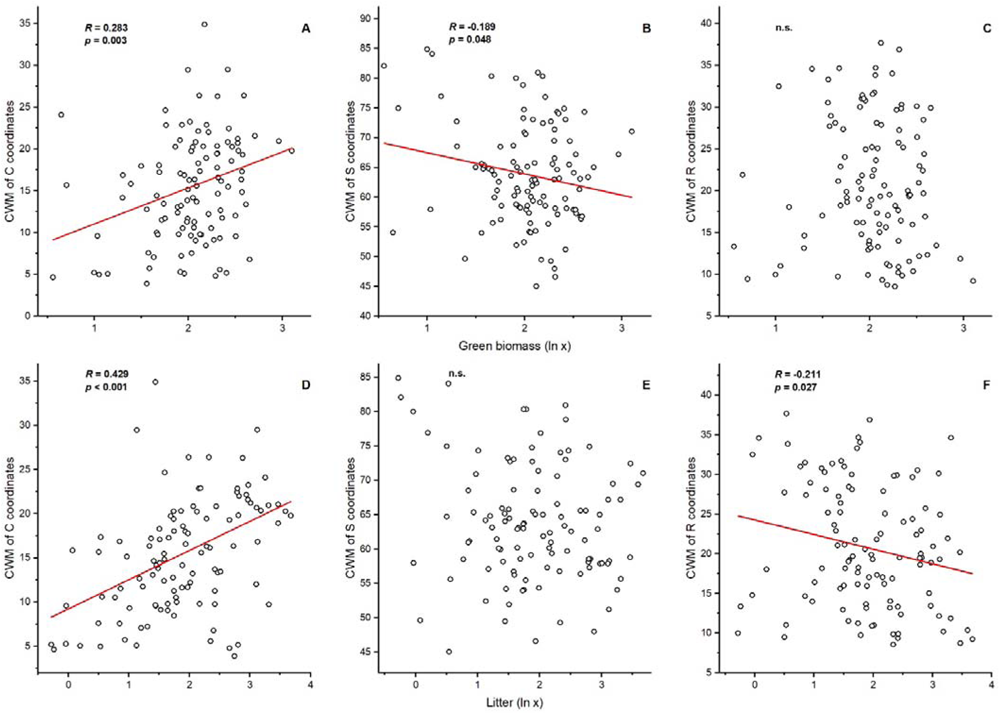
Correlation between green biomass and litter and CSR coordinates. Correlations were calculated by Pearson’s product moment. Notations: CWM = community-weighted means, n.s. = non-significant. Biomass scores are in grams for the 20 cm × 20 cm plots.

## 4. Discussion

### 4.1. Biomass and species richness of the sampled communities

We hypothesised that the biomass and species richness scores and the species composition of the sampled communities along the sampled gradients will be significantly different. These were supported by our results. The highest species richness, evenness and Shannon diversity values were found in loess grasslands. This is well in accordance with the findings of Kelemen et al. (2013), who analysed the relationship between biomass production and species richness. It was found that with intermediate levels of biomass production the loess grassland has the highest species richness compared to other grassland community types in alkaline landscapes. The lowest green biomass and litter was found in short-grass steppes, which is also in line with the former findings of Kelemen et al. (2013). In the sense of Grime (1977), stress is defined as those factors that prevent biomass production. In line with this, the highest levels of stress can be expected in sites with the lowest amount of produced biomass – which latter is in short-grass steppes. Beside of the salinity stress – which is rather evident in this grassland community type – there is also a high level of nutrient stress (Kelemen et al., 2013). We also found that a relatively high amount of cryptogam biomass (including various stress-tolerant lichens and nitrogen-fixing terrestrial cyanobacterial group, *Nostoc*) was typical in these communities. A neutral to negative effect of *Nostoc* species on the germination and early species establishment of alkaline plant species was validated by Sonkoly et al. (2017) except for *Beckmannia eruciformis*; which might have an effect on species composition.

The highest level of litter accumulation was found in alkaline meadows, which are characterised by the high dominance of several graminoids including *Alopecurus pratensis*, *Agrostis stolonifera* and *Elymus repens* (see Figure 5). It was found formerly that the amount of litter shows strong correlation with the biomass production of dominant graminoids (Facelli and Pickett, 1991; Kelemen et al., 2015). Litter also has mulching effect, which supports the retainment of soil humidity, decreases direct light irradiance, and mitigates the temperature extremities of the soil surface (Grace and Pugesek, 1997; Foster and Gross, 1998; Xiong and Nilsson, 1999; Eckstein and Donath, 2005). However, this higher magnitude of litter accumulation is also responsible for the lower species richness detected in these grassland community types of as litter also suppresses the germination of light-demanding gap strategist plants (Ruprecht et al., 2010). Surprisingly, the green biomass was similarly high in the loess grasslands, however, the litter accumulation was much lower than in alkaline meadows. This can be explained by the fact that loess grasslands harbour a high diversity and biomass of forbs, which are not likely to contribute to litter accumulation in the same magnitude as tall graminoids in alkaline meadows. Also, tussock-forming graminoids like fescue species do not form such a dense canopy cover as clonally spreading grasses, leaving gaps for the establishment of forb seedlings (Kelemen et al., 2015), which leads to a higher species evenness detected for the loess grasslands compared to other grasslands in this study.

### 4.2. The CSR strategy spectra of the sampled communities

We found high levels of stress selection in all studied communities (>50% of the community weighted means of S coordinates in all studied communities). Our results are in accordance with the findings of Negreiros et al. (2014), who found that the stress-tolerant strategy was predominant in grassland communities under harsh environmental conditions. The high degree of S selection is clearly visible in Figure 7 and 8, where the points representing the biomass samples of the studied communities are located close to the bottom-right corner of the ternary plot. We hypothesised that the sampled communities will be separated from each other based on the CSR strategy spectrum. This hypothesis has not been validated by the results as the studied communities were not separated in the ternary diagram generated using community-weighted coordinates of CSR axes.

One possible explanation for the lack of separation of the communities based on CSR strategies is that the applied classification system performs well if we compare contrasting regions or habitat types, but it seems to be not sensitive enough to compare communities with a limited and highly overlapping species pool. While the CWMs of C and R coordinates were significantly different between the studied communities, we found no significant differences between the S coordinates (Figure 6). The CSR classification is designed to quantify the magnitude of stress and disturbance in the habitats or communities under consideration, not to differentiate between the natural or the temporal patterns of these. Plants are facing several stress factors in alkaline habitats. These factors include (1) high osmotic pressure caused by accumulated salts, (2) ion toxicity and an unbalanced ion concentration, (3) unfavourable soil conditions in terms of compactness, water capacity and soil structure, (4) suboptimal soil pH caused by alkaline soils (mostly caused by natrium and potassium carbonates and sulphates), and (5) water and nutrient deficiency (Füzy et al., 2010). In the sampled grasslands, the characteristic species must tolerate high levels of stress at least in a particular period of the year, and also its spatiotemporal patterns and magnitude because of the high patchiness of the vegetation and variations in microtopographic differences is highly variable even within a particular community (Valkó et al., 2014; Deák et al., 2015; Matinzadeh et al., 2022). In loess grasslands, the plant community is shaped mainly by drought stress, while the nutrient availability in the typical chernozemic soils of loess grasslands is generally not limited and the salt content of the soil is in general low (Valkó et al., 2021). These favourable conditions enable high levels of biomass production and competition. Alkaline meadows, the deepest-lying communities we sampled, are covered by shallow water until late July – early August in most years. While salinity is relatively high, the salt content is in a solution during most of the year, and typically high in lower soil layers. After the community dries out in the summer, we can expect high levels of drought stress. The third grassland community type represents a very interesting situation. While it is typically inundated until early spring, it then dries out and most of the salt accumulates in the upper soil horizon, which creates conditions affected by both salt and drought stress. The nutrient contents of the soils formed here are rather low, causing nutrient stress as well (Valkó et al., 2014). Grime defined stress in his CSR framework as factors that prevent biomass production (Grime, 1977). Based on this assumption, the community with the lowest amount of biomass production can be exposed to the highest levels of stress. We found the lowest amount of green biomass in the short-grass steppes. Despite this, the CSR strategy spectra of the studied communities was not separated. The presented analysis of CSR strategy spectrum considers the biomass proportions of species and does not consider the total amount of biomass in the studied communities. Integrating the total amount of biomass into CSR spectrum analyses may provide a better insight into stress levels in the studied communities.

Selective grazing by livestock may be another reason for the lack of separation of the communities based on the CSR strategy system. The study sites were lightly to moderately grazed by beef cattle in the vegetation period, which can strongly influence the species and trait composition. It was found by several former studies in the region that grazing, especially differences in grazing intensity can fundamentally shape species composition and CWM trait values and that grazers are often selective for particular species and/or for community biomass (Tóth et al., 2018; Török et al., 2018). 2022 was a very dry year in the region with an especially dry spring, which probably strongly influenced the early germination and establishment of short-lived species, characteristic in alkaline and loess grasslands. Drought most likely affected the sampled communities unevenly, and its effects were probably the most pronounced in communities regularly affected by drought stress even in typical years. This might also be reflected in the observed CSR strategy spectrum of loess grasslands, which is affected by the high proportion of perennial species in their biomass samples and by their effects on the CSR coordinates. Finally, while we used regionally measured data for traits (Sonkoly et al., 2023), some authors argued that in trait-based studies, on-site measurements of traits would be the most appropriate (Cordlandwehr et al., 2013). For most species we do not know the intraspecific trait variability, but we can assume that it can be considerable for species characteristic in multiple grassland community types in alkaline habitat complexes with various magnitudes and types of stress factors.

## 5. Conclusions

Based on the results we can conclude that the current CSR classification is too robust to delineate grassland communities in alkaline landscapes which are markedly different in composition. We point out that the CSR classification system is based on the magnitude of stress, and it is not able to differentiate between the various forms of stress which might be responsible for the marked compositional differences. Therefore, developing an index focusing specifically on the functional characterisation of stressed communities and able to differentiate between different forms of stress (e.g., water and/or salinity stress) would be highly beneficial for future studies. To obtain the broadest possible view of the CSR strategy spectrum of different grassland communities in alkaline landscapes, it would also be necessary to sample in years with markedly different weather conditions and in stands under differing management regimes.

## Acknowledgements

The work was supported by the ÚNKP-23-3-I. New National Excellence Program of the Ministry for Culture and Innovation from the source of the National Research, Development and Innovation Found. The authors were supported by the National Research, Development and Innovation Office [PT: NKFIH KKP 144068, K 137573; JS: PD 137747] during the manuscript preparation. The work of JS was supported by the Bolyai János Scholarship of the Hungarian Academy of Sciences [BO/00587/23/8]. This project has received funding from the HUN-REN Hungarian Research Network. The suggestions of László Erdős to the earlier version of the manuscript were gratefully acknowledged.

